# Genetic loss of the ubiquitin ligase RNF126, an AID interacting partner and modifier, affects strand targeting during somatic hypermutation of antibody genes

**DOI:** 10.1101/469775

**Authors:** Nicholas Economos, Rebecca K Delker, Pete Stavropoulos, F. Nina Papavasiliou

## Abstract

Activation-induced cytidine deaminase (AID) initiates somatic hypermutation (SHM) and class switch recombination (CSR) in B lymphocytes by catalyzing the introduction of deoxyuracil: deoxyguanine mismatches into the DNA of the transcribed Ig locus. Repair pathways then process these mismatches to produce point mutations in the Ig variable region or double-stranded DNA breaks in the switch region followed by deletional recombination. It has been suggested that post-translational modifications on AID mediate a number of these different decisions, ranging from global targeting (Ig vs the genome), local targeting (variable vs switch region; transcribed vs non-transcribed strand) as well as process-appropriate DNA repair. Here we demonstrate that absence of RNF126, an E3 ligase shown to mono-ubiquitylate AID, results in a specific strand targeting defect in SHM, producing substantial G>C bias; strickingly, loss of RNF126 was also associated with tandem indels within the variable region (JH4 intron) but only a slight increase in the types of chromosomal translocations that are characteristic of deregulated AID. Conversely, these findings suggest that mono-ubiquitination of AID, likely *in situ*, is necessary for the proper removal of the protein from the non-transcribed strand, thus producing both optimal patterns of SHM and also limiting the number of indels within the target locus.

## Introduction

Antibodies are initially assembled in the bone marrow in an antigen-independent manner by site-specific recombination (mediated by the RAG1/2 recombinase) and are further diversified in the secondary lymphoid organs through somatic hypermutation (SHM) and class switch recombination (CSR). SHM alters antibody affinity by introducing nucleotide changes in the antigen-binding variable region of antibodies. B cells producing antibodies with improved antigen affinity are positively selected during the process of affinity maturation ^1^. CSR is a region-specific recombination reaction that replaces one antibody-constant region with another, thereby altering antibody effector function while leaving the variable region and its antigen binding specificity intact ^1^. While CSR and SHM are very different reactions, both are initiated by activation-induced cytidine deaminase (AID) ^2,3^, which introduces programmed DNA damage (a deoxyuracil-deoxyguanine (dU:dG) mismatch) at the Ig locus ^4^.

Even though the vast majority of AID activity is sequestered to the Ig loci, AID can also induce “off-target” DNA damage, including point mutations in oncogenes such as *bcl6* ^5^, as well as double-stranded breaks that result in oncogenic chromosome translocations such as those between c-myc and IgH (c-myc/IgH) ^6^. Thus, maintaining genomic integrity requires strict control of AID activity and localization; this is achieved at transcriptional, post-transcriptional as well as post-translational levels.

Post-translationally, AID activity is regulated by phosphorylation and ubiquitination ^7,8^. To date, biochemical experiments have demonstrated that phosphorylation of certain residues (e.g. at Ser38 or Thr140) positively regulates AID activity whereas phosphorylation of others (e.g. Ser3) act to suppress AID activity. However, most of these modifications are initially assessed with regard to their effect on CSR; their effect on SHM is deduced from experimental systems (DT40 gene conversion, mutation in 3T3-NTZ fibroblasts) that are not fully faithful to hypermutation at the Ig locus ^8^.

Here, we focus on AID mono-ubiquitylation, which was first observed in nuclei of mutating cell lines (BL2) as well as nuclei of switching cells ^7^. Using a protein-protein interaction screen, we had previously isolated the E3 ligase RNF126 and demonstrated that it selectively modified (mono-ubiquitylated) AID *in vitro* and *in vivo* (in HEK293T cells) ^9^. To assess the effect of AID ubiquitylation within the relevant cell types, we examined aspects of CSR and SHM in B cells that were generated to conditionally lack the RNF126 E3 ligase. Our experiments demonstrate subtle but specific defects in the strand targeting of SHM in the absence of the modifying enzyme.

## Results

### Depletion of RNF126 from CH12F3 cells results in reduced CSR

We have recently discovered that the RING E3 ligase RNF126, with E2 UbcH5b, can mono-ubiquitinate AID in cell-free assay conditions and in 293T cells ^9^. To begin to assess the functional implications of RNF126-mediated AID ubiquitination, we turned to the CH12F3 cell line, an easily accessible model of CSR, thought to faithfully recapitulate the reaction ^10^. CH12F3 cells express surface IgM, and can switch to surface IgA upon induction with proper stimulation (CD40, IL-4 and TGF-beta - ^10^). They can also be infected with lentiviruses carrying shRNA hairpins: transfection of an shRNA that knocks down levels of AID protein leads to a near-complete ablation of CSR under these conditions (Fig. 1A and 1B). Transfection of shRNAs targeted to RNF126 result in diminished frequencies of CSR (Fig. 1A and 1B), commensurate with the levels of RNF126 protein remaining (Fig. 1C). Hence, an acute knockdown of RNF126 from the CH12F3 cell line leads to a substantial reduction of switching from IgM to IgA (∼70% reduction from WT levels, comparable to what has been reported for *spt5* ^11^ or *ptbp2* ^12^).

**Figure 1.**
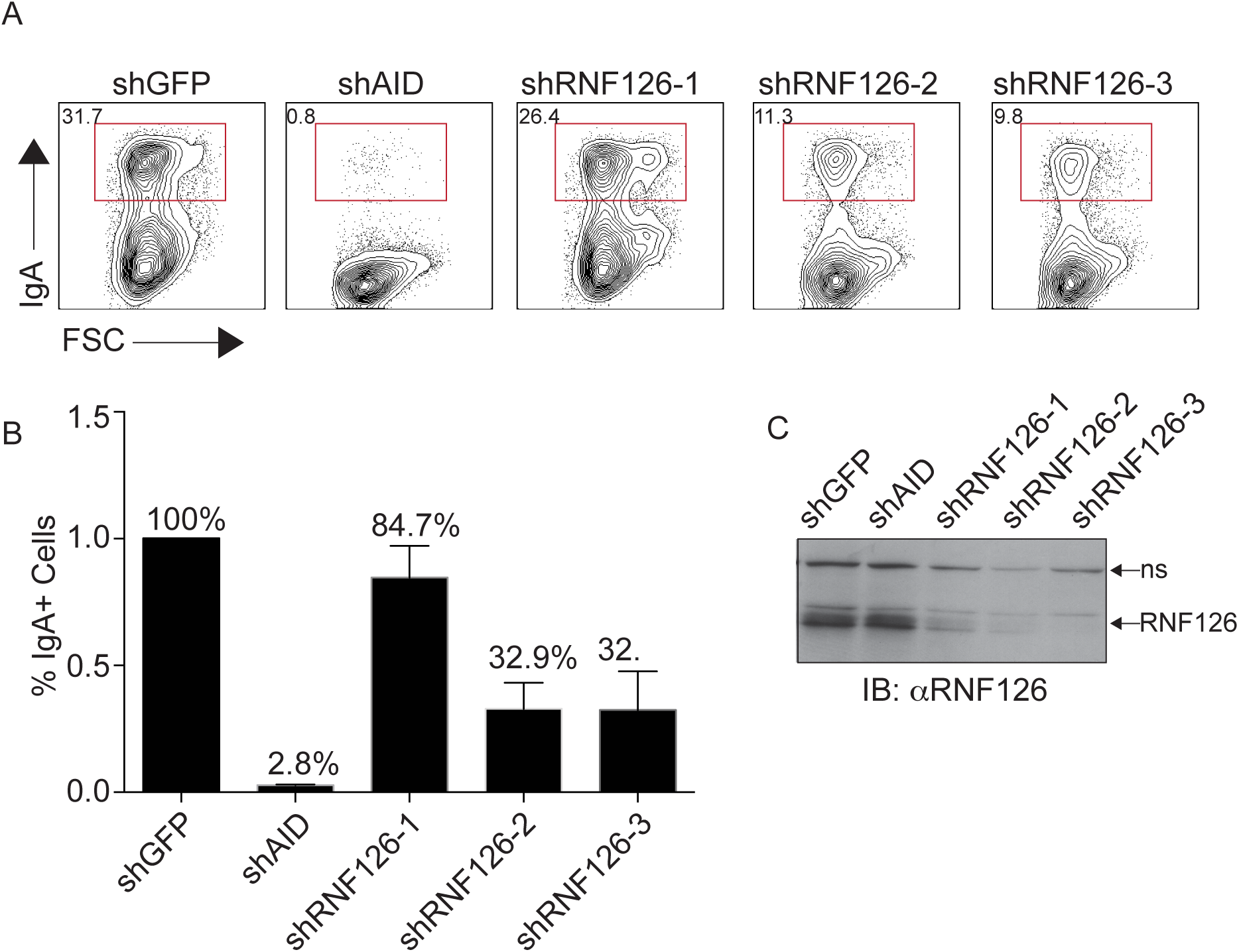
Silencing of RNF126 from CH12F3 B cells leads to reductions in CSR levels. CH12F3 cells were infected with retroviruses expressing siRNA hairpins to GFP (negative control), AID (positive control) and RNF126 (three hairpins). After selection, infectants were stimulated to switch from IgM to IgA using a cytokine cocktail (CD40L, IL4 and TGFb). (**A**) Representative FACS plots and (**B**) agreggate data from 3 different experiments are shown. (**C**) Western blot of cell lysates (normalized for protein amounts) were probed with anti-RNF126 antibody to assess depletion. siRNA hairpins to RNF126 show variable reductions in protein levels, that correlate with reductions in CSR.

### Conditional deletion of RNF126 from B cells results in mildly reduced CSR levels

RNF126 is a relatively uncharacterized gene, and a full knock-out resulted in perinatal lethality and substantially skewed mendelian ratios (Fig. S2A and not shown). We thus generated an *rnf126^-/-^* conditional knock-out strain (Fig. S1) which was crossed to mb1-Cre to achieve deletion of RNF126 early in B cell development ^13^. Loss of RNF126 from the early B cell compartment onward, did not appear to affect B cell development in the bone marrow or levels of B cells in the spleen (Fig. S2).

We then sought to determine if primary B cells derived from RNF126^Fl/Fl^ mb1^Cre/+^ mice displayed compromised levels of CSR. Specifically, we isolated splenocytes from these mice and stimulated them with IL-4/aCD-40 to induce a switch from IgM to IgG1. In comparison to wildtype littermates, RNF126-deficient B cells are able to undergo CSR, but do so at consistently reduced frequencies, at every time point post stimulation (Fig. 2A-B and Fig. 2D). Recently, RNF126 has been proposed to support cancer cell proliferation by targeting p21 in cancer cell lines ^14^ and siRNA-mediated reduction of RNF126 levels was shown to result in loss of viability. Aside from a mild alteration in proliferation kinetics however (Fig. 2C), we did not observe increased cell death in these cultures (not shown). Thus, the loss of viability phenotype others have reported, might be restricted to tumour settings.

**Figure 2.**
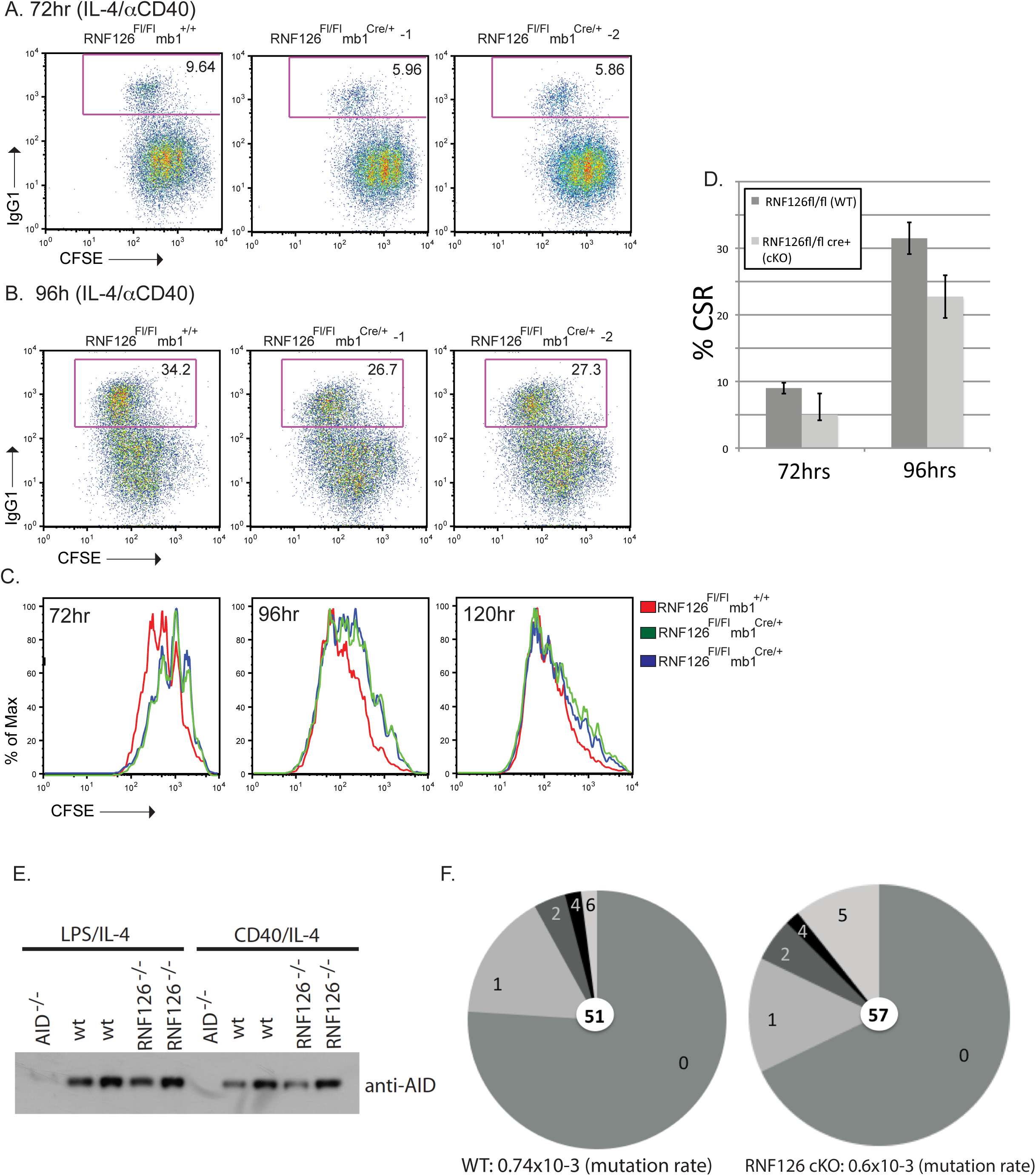
Genetic loss of RNF126 affects CSR levels. Naïve splenic B cells derived from RNF126^Fl/Fl^ mb1^+/+^ and RNF126^Fl/Fl^ mb1^Cre/+^ mice were stimulated *in vitro* with aCD-40 and IL-4 to induce a CSR event to IgG1. FACS analysis completed at 72hr and 96hr post stimulation was used to determine the percentage of cells which express surface IgG1. (**A-B**) Representative FACS plots (**C**) Cell proliferation assessed by CFSE staining (**D**) Histograms depicting slight reductions in IgG1 levels at 72h and 96h after stimulation (8 paired biological replicates per condition). (**E**) Bulk levels of AID do not vary between mice (lysates normalized as explained in materials and methods).(**F**) Pie charts of clonal mutations recovered from the Smu core region (a measure of AID occupancy) were determined as in ^40^. Mutation rates were determined using unique mutations only. P-values, determined by a 2-tailed Fisher’s exact test, are shown.

AID protein is highly regulated at multiple levels and a mild reduction of CSR such as we observe (aggregated in Fig. 2D) could result from alterations in protein amounts, subcellular localization or sub-nuclear targeting ^1^. RNF126-deficient B cells do not display alterations in AID levels (in bulk – Fig. 2E), and within these cells, AID can certainly enter the nucleus, and deaminate the switch region generating characteristic levels of Smu mutations (Fig. 2F), leading to initiation of appropriate DNA repair and resulting in CSR with near-wildtype kinetics.

### Conditional deletion of RNF126 from B cells affects the processivity of AID on the Ig locus

CSR and SHM are both initiated by AID catalyzed deamination, yet the differences between them are substantial. Unlike CSR, an *ex vivo* or *in vitro* system that can faithfully recapitulate SHM does not exist currently and SHM is usually assessed by isolating B cells from germinal centers, and cloning and sequencing an intron within the Ig locus that is not the subject of selection (usually the JH4 intron).

To assess whether RNF126 deficiency affected SHM, we isolated CD19+GL7+Fas+ germinal center B cells from Peyers’ patches excised from aged (9mo old) RNF126^Fl/Fl^ mb1^Cre/+^ mice. From this material, we cloned and sequenced the JH4 intron and compared wildtype and RNF126-deficient cells for a number of SHM-related variables, such as the local base targeting preference of AID within the Ig locus, the strand biased targeting or repair of AID-mediated lesions, and ultimately, the efficiency of SHM (both in terms of frequency overall, and in terms of mutations per clone).

We found that mutation frequencies were overall slightly lower in the RNF126-/- cells compared to their wildtype littermates (1 vs 1.5%). However, from RNF126-/- B cells we recovered a larger number of clones that were highly mutated (>10 mutations) as compared to the wildtype (Figure 3A, p=0.008; one-tailed Fisher’s exact test). This was both true for JH4 regions amplified from GC B cells isolated from Peyer’s patches, and for JH4 regions amplified from GC B cells after immunization with NP-CGG (supplemental Fig. 3). These findings imply that when RNF126 is not present, unmodified AID stays on the locus longer.

**Figure 3.**
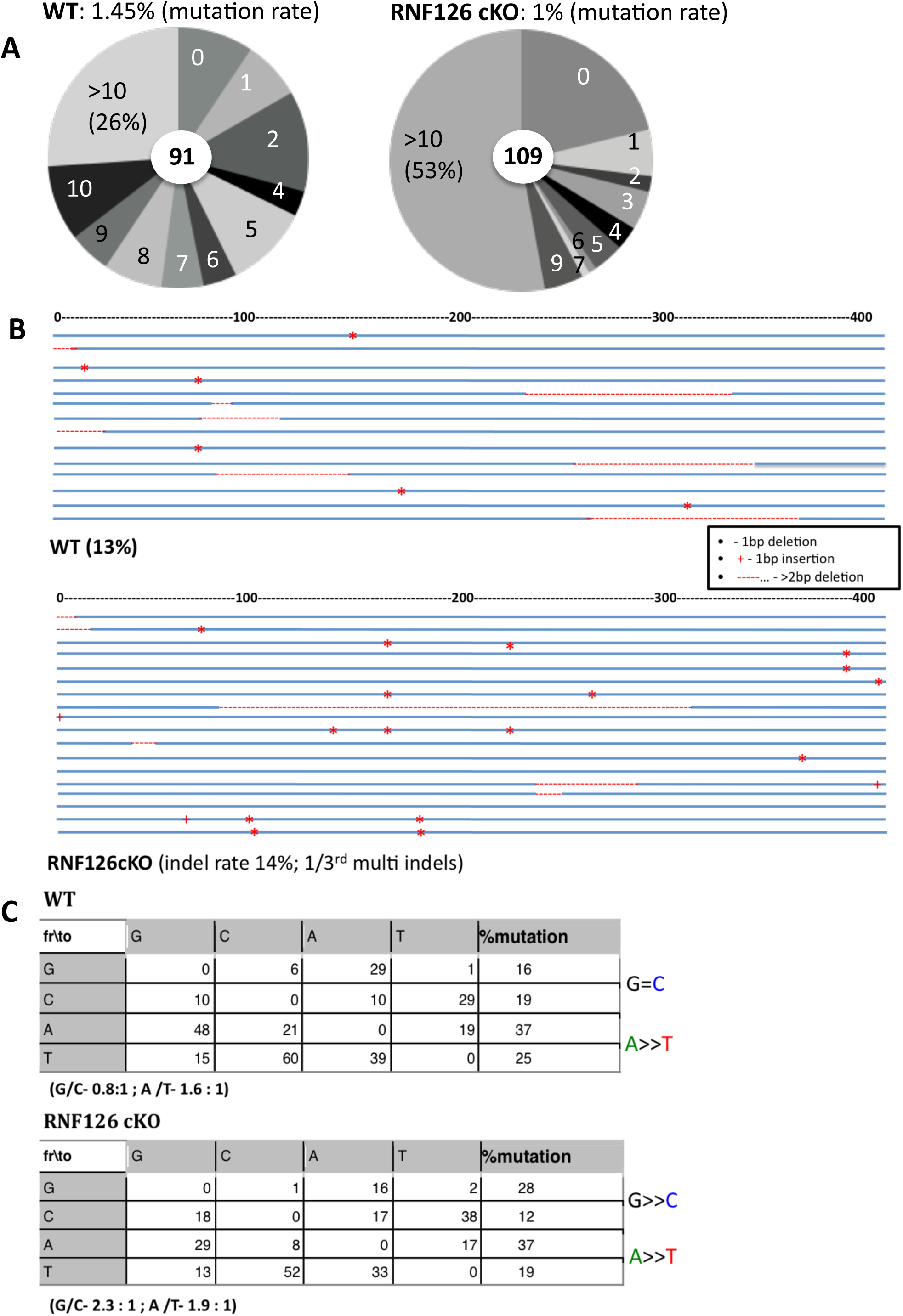
Genetic loss of RNF126 affects rates and patterns of SHM. To assess SHM in the JH4 intron we sorted CD19+GL7+Fas+ germinal center B cells from Peyer’s patches collected from aged (9mo) RNF126^Fl/Fl^ mb1^+/+^ and RNF126^Fl/Fl^ mb1^Cre/+^ mice. We then amplified the JH4 region using genomic DNA from sorted cells. (**A**). Pie charts show the fraction of total sequences that contain the indicated number of mutations for the two genotypes. Mutation rates are depicted at the top of each chart. (**B**) Sequences with indels that are characteristic of SHM are shown as line diagrams (as per the legend on the right). Rates of indels are depicted at the bottom of each set of line diagrams. (**C**). Mutation tables depict the % of mutation from each base in the sequence to every other possible base. Values are corrected for base composition, and GC/AT ratios are shown at the bottom of each table (with mutations normally demonstrating strand bias in A/T ratios but balanced G/C).

SHM has also been associated with a low but measurable rate of insertions and deletions which account for 5-10% of all events, depending on the study ^15^. Although usually quite short, indels can be of a variety of sizes ^16^, and are thought to be the outcome of point mutations on opposite strands that occur sufficiently closely in space and time that they result in a DNA double strand break. They are also thought to be proximal measures of AID activity: AID overexpression leads to an increase in indel rates from about 5% to 25% ^17^. We therefore sought to measure the numbers of clones with indels in our experiments, and found that the rates are very similar between RNF126-deficient animals and their wildtype littermates (∼8%). This supports the notion that the encounter of AID with the Ig locus is not perturbed in the RNF126-/- B cells. But whereas the frequency of clones with indels is not different between the strains, the nature of indels, is. Specifically, whereas in wildtype B cells clones with indels contain a single indel event (confirming prior studies – ^16,17^), we recovered a number clones with more than one, and sometimes up to three distinct indels per clone (accounting for 30% of clones with indels in the conditional knockout cells - Fig. 3B). Together with the increase in numbers of mutations per mutated clone we noted in RNF126-/- B cells (Fig. 3A), this finding suggests a higher half life of unmodified AID on the locus. Specifically, a presumed loss of ability to be removed from DNA in a timely fashion, would be predicted to impact the apparent processivity of AID (both with regard to mutations per clone and with regard to the sequential indels in mutated clones) such as we observe.

### Conditional deletion of RNF126 from B cells results in altered patterns of SHM

AID displays local sequence preference for certain "hotspot" motifs in DNA, and because this preference is substantially similar *in vivo* and *in vitro*, it is thought that it is an inherent biochemical attribute of the molecule. This preference in hotspot targeting (e.g preference to deaminate the C residues in WRC (W = A or T, R = A or G) hotspot motifs) was not altered between wildtype and RNF126 deficient cells (not shown).

*In vivo*, the rate of mutation recorded from the non-transcribed strand, is equal from C or G (a C on the transcribed strand), suggesting that AID deaminates both substrate DNA strands equally well, during SHM ^18,19^. Indeed, it appears that AID induces dUs equally on both strands of the DNA ^20^. In contrast to the equivalence of mutations from C or G, that are taken as evidence for balanced AID targeting, adenine (A) bases are mutated twice as frequently as the complementary thymine (T) bases. This has been directly linked to the type of DNA repair recruited to the locus, and specifically, to the function of the TLS polymerase pol-eta, a low fidelity polymerase that preferentially synthesizes mispairs when copying T bases located on the transcribed strand ^21,22^, together with a number of DNA repair proteins that are normally part of the error-free mismatch repair complex ^23^.

The A>T bias was largely intact in the RNF126-deficient B cells, suggesting that the presumed lack of AID modification did not alter the recruitment of the “canonical” error-prone repair pathway that spreads mutation through the locus (Fig. 3C). Surprisingly however, we found a strong bias in mutations from C or G as recorded from the non-transcribed strand: that is, Cs on the template strand are mutated more frequently. Whereas in the wildtype SHM G=C, SHM in RNF126-/- B cells show a substantial preference for G>C (Fig.3C). This finding is recapitulated in data from JH4 amplification and sequencing deriving from germinal center B cells, which were sorted after immunization with NP-CGG (supplemental Fig. 3).

Together with the effects caused by the absence of RNF126 on AID processivity and on strand bias, aspects of the reaction that are thought to be proximal to the introduction of dU in DNA, these experiments suggest that RNF126-mediated modification of a factor very close to the initiation of the introduction of dU in DNA (likely AID) is necessary for optimal SHM.

### Conditional deletion of RNF126 from B cells slightly increases the level of AID-mediated translocations from the Ig locus

Increasing the time of occupancy of AID on the Ig locus might be predicted to result in the types of reciprocal chromosomal translocations that, while rare in wildtype cells, are the hallmarks of a number of B cell tumours, most prominently B cell lymphomas. To test for this, we queried RNF126-/- cells for reciprocal translocations between the IgH and c-myc loci (Fig. 4A), which occur between a rate of ∼0.5 in 10^6^ in the wildtype and which are substantially increased in cells that ectopically overexpress AID (∼50 in 10^6^) ^24^ or cells that are deficient in DNA repair (e.g. pol-Q; ∼3 in 10^6^) ^25^. We compared B cells from mice that were RNF126-/- (3 biological replicates) to their wildtype littermates (4 biological replicates) and to cells mice that were doubly deficient (ie AID-/-RNF126-/-; 4 biological replicates). These cells were stimulated to undergo CSR, and were collected and processed for translocation assays 72hrs after stimulation. Doubly deficient cells showed no translocations whatsoever, of any type (data not shown), much as has been reported for AID deficient cells. We recovered one reciprocal translocation in ∼4 million wildtype cells, on average, in all replicates (∼0.25 in 10^6^). In contrast, we recovered ∼11 reciprocal translocations from 6 million RNF126-/- B cells, on average, in all replicates (∼1.8 in 10^6^). Examples of translocation amplicons where the Ig locus (chromosome 12) is translocating to c-myc on chromosome 15, are shown in Fig. 4B; the quantification of multiple such examples, is in Fig. 4C). Because AID is not overexpressed in these cells, which are also repair proficient, we conclude that the slightly increased translocation rates here reflect increased occupancy of AID on the bottom (transcribed) strand of the Ig locus (also reflected in the processivity of both mutations and indels that we observe) and the DNA repair pathways that are called upon to remove it.

**Figure 4.**
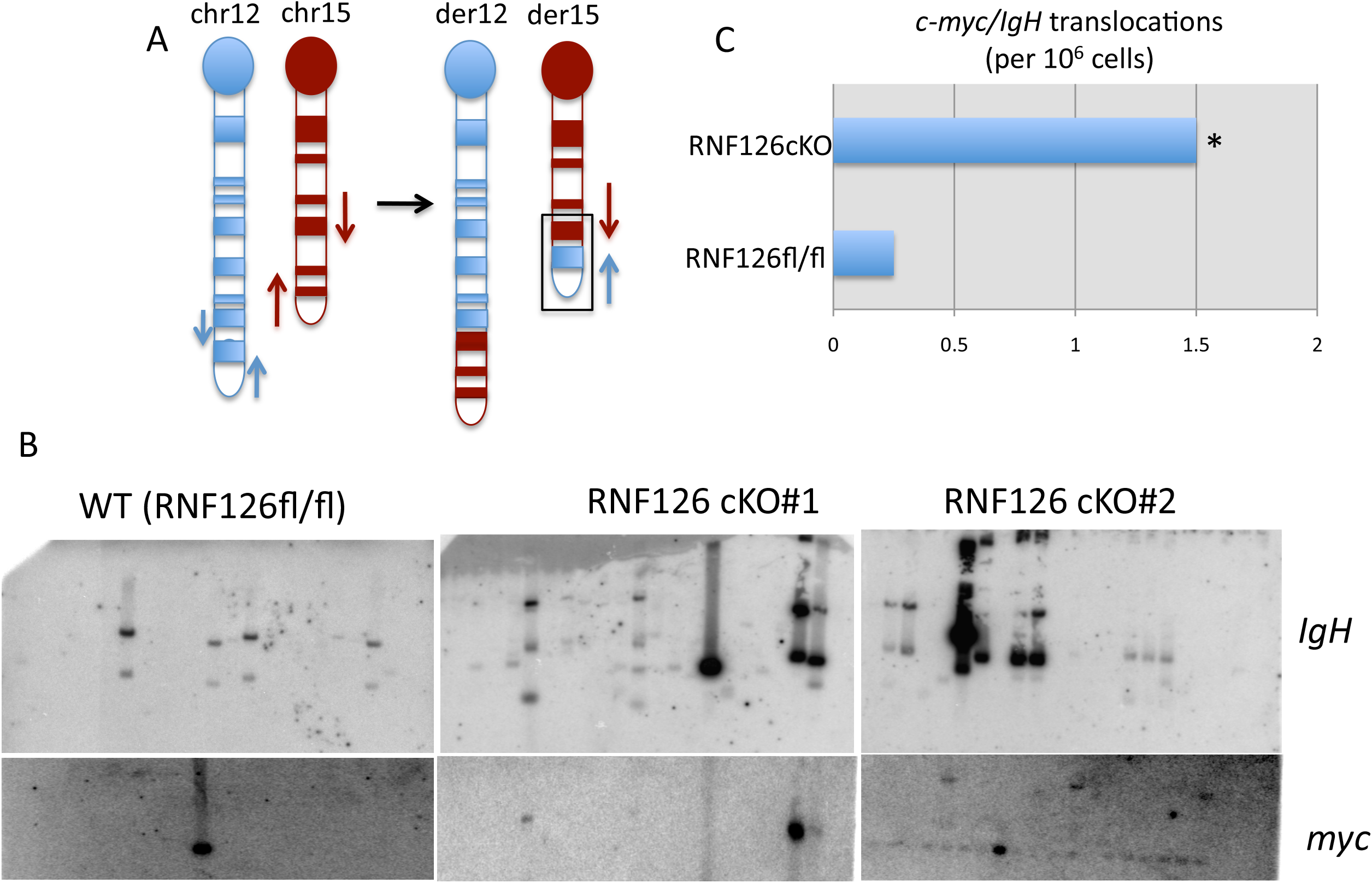
Genetic loss of RNF126 leads to a slight but not significant increase in chromosomal translocations. (**A**) Schematic representation of PCR assay for cmyc/IgH reciprocal translocations. Primers used to detect translocations are represented as gray arrows. (**B**) Representative southern blots showing c-myc/IgH translocations in RNF126^Fl/Fl^ mb1^+/+^ (WT) and RNF126^Fl/Fl^ mb1^Cre/+^ (RNF126cKO) B cells, 72hrs after stimulation to undergo CSR (probed with c-myc or IgH oligo probes, as indicated). Each lane represents genomic DNA from ∼100,000 cells. c-myc/IgH translocations. (**C**) Frequency of translocations obtained from RNF126^Fl/Fl^ mb1^+/+^ (WT) and RNF126^Fl/Fl^ mb1^Cre/+^ (RNF126cKO) B cells (after analyzing 4 biological replicates per genotype, 8x10^6^ total per replicate, and counting only reciprocal translocations, ie positive for both IgH and myc probes). The p=0.6 value was determined with chi-squared test with Yates correction for large values.

## Discussion

The single-strand DNA-specific cytidine deaminase activation-induced cytidine deaminase (AID) is essential for antibody diversification both through CSR and through SHM ^2,3^. AID deaminates cytosines within the transcribed Ig loci, and the deaminated DNA engages proteins of ubiquitous repair pathways to generate DSBs (in the case of CSR) or to spread the mutation from the original dU:dG base pair to the surrounding area, creating stereotyped patterns of SHM that are required for efficient diversification of the antibody binding pocket ^26^.

All aspects of AID biology are highly regulated, from its subcellular localization (cytoplasmic retention vs nuclear import and export), to its targeted deposition onto the Ig locus once in the nucleus ^27^. Once at the Ig locus, AID is further targeted to the VDJ exon (for SHM) or the S region intron (for CSR) where it acts on either strand to generate programmed DNA damage in the form of deaminated cytosine, which then differentially resolved either into stereotypical mutational patterns that spread well beyond the original dU:dG mispair (as is the case for SHM) or into DNA DSBs (in the case of CSR). Conversely, mistargeting of AID activity or the resolution of AID lesions to non-Ig genes has been implicated in chromosomal translocations and pathogenesis of B cell lymphomas ^28^. But despite intense efforts and a number of recent advances, the exact mechanism of AID targeting to the Ig, as well as the mechanisms by which it differentially attracts DNA repair factors, remain unclear.

AID is a relatively small protein (25kDa) so it was originally hypothesized that much of the regulation discussed above is mediated by post-translational modifications. Indeed, AID is heavily modified: for example, it is phosphorylated at multiple residues ^8,29^ and patterns of phosphorylation might be important to determine some of these functions (for example the absence of phosphorylation at specific residues can differentially affect SHM vs CSR ^8^). Here we have focused on AID ubiquitylation as mediated by the E3 ligase RNF126. We have previously shown that *in vitro* and in overexpression systems, RNF126 does mono-ubiquitylate AID with some selectivity ^9^. *In vivo*, absence of RNF126 from B cells, especially in the context of SHM, results in increased processivity of both mutations and indels, and in differences in strand bias. These however do not result in significant increases in the rates of translocation between the Ig and myc loci, suggesting that the effect we see on SHM in the absence of RNF126 are limited to AID on the Ig locus. Focusing on the mutation and indel data, these suggest either that the bottom strand is targeted at increased rates (which then ought to result in alterations in the spreading of mutation to neighboring A/T bases, <u>unlike</u> what we observe) or, that AID must be actively removed from DNA, and that optimally, active removal from the locus requires modification *in situ*, either prior to, or concurrently with deamination of dC to dU.

The introduction of strand-balanced, AID-generated dU in DNA can result from at least three mechanisms. First, a biophysical, movement-related mechanism where negative supercoiling is generated on DNA on the wake of the RNA polymerase, dynamically generating short stretches of quasi-stable ssDNA on both strands ^30,31^. Second, a mechanism that requires stalling of a transcription complex, with AID directly attacking the non-transcribed strand as well as the template strand after the exosome-mediated “peeling” of RNA off the latter (this is likely the predominant mechanism in CSR - ^32^). Third, a transcription-complex-directed mechanism, where AID directly attacks the non-template ssDNA presented on the RNA polymerase during transcription, while attack on the template strand DNA requires a subsequent antisense transcription event ^33^. Of these, the latter would predict that the AID-mediated initial deamination of dC to dU on both strands happens through an identical mechanism (attack on the non-template strand) but during two distinct waves of transcription: through deamination at non-template dC, with subsequent deamination at non-template dC on the opposite strand (recorded as dG on the top strand). Thus, the overall equivalence of mutations from C or G, would not necessarily imply that both strands are handled similarly, regarding the resolution of dU:dG into a mutation.

*In vitro*, AID displays remarkably high affinity for single-stranded DNA as indicated by the low dissociation constants and long half-life of complex dissociation ^34^, and these biochemical findings have been taken to suggest that AID protein may persist the Ig locus after deamination, possibly acting as a scaffolding protein to recruit other factors. Given our *in vivo* data, we wondered whether the G>C strand imbalance we see, might be reflective of equivalent targeting of both strands, but an inability to properly remove AID from the template strand, unless it is modified. Essentially, we wondered whether ubiquitylation might be a “handle” with which a “mutasome” complex might remove AID from the locus, to allow for timely DNA repair so that transcription can proceed. In the absence of ubiquitylated AID then, the cell might resort to a salvage pathway of AID removal through DNA repair mechanisms that normally deal with DNA:protein adducts. Such mechanisms would include CtIP involvement, or involvement of the Mre11-Rad51-Nbs1 (MRN) complex. Both of these have been shown to act on the locus ^35,36^, and the MRN complex has been specifically associated with repair of template strand C ^37^.

In contrast to SHM, CSR in RNF126-/- B cells is only mildly impacted (Figure 2). This is either because targeting of AID to the switch or variable regions happens through substantially different mechanics (e.g. semi-stable R-loop formation vs transient single stranded regions), or, because the targeted removal of AID from switch regions, such as we suggest, though still necessary for transcription to proceed, is not essential for every single binding event (given that the regions will be excised from the genome).

We have not, however, succeeded in demonstrating that ubiquitylated AID is present within the chromatin of the Ig locus. This is likely for technical reasons: AID levels are not high in GC B cells, most of the protein (∼95%) resides in the cytoplasm and even within the nuclear fraction, a small amount can be found on chromatin ^38^. Coupled with the fact that existing anti-AID antibodies are suboptimal in their ability to immunoprecipitate and/or detect AID, and that anti-Ub antibodies are usually not able to detect mono-ubiquitylated protein species, at present, we cannot formally demonstrate that in the wildtype, AID is monoubiquitylated by RNF126 *in situ*. However, taken together, our data do suggest that monoubiquitylation of AID is necessary for optimal removal of the protein from DNA, allowing DNA repair factors to access the dU-containing strand in a timely fashion so that transcription can proceed.

## Acknowledgements

This work was supported by National Institutes of Health Grant #CA098495 (National Cancer Institute) (to FNP), Starr Cancer Consortium grant# I4-A447 (to FNP), by graduate fellowships from the National Defense Science and Engineering Foundation and the National Science Foundation (to RKD) and by a postdoctoral fellowship from the Beckman Foundation (to PS).

## Contributions

The RNF126 conditional mouse was originally generated and initially characterized by RKD, who also generated all the supplemental figures as well as figure 1, and contributed data to figure 2. PS and NE contributed data to figure 2 and generated data for figures 3 and 4. All authors had input in the final manuscript.

## Materials and Methods

### Generation of an RNF126 conditional knock-out mouse model

ES cells containing the targeted RNF126 allele were purchased from the EUCOMM International Knockout Mouse Consortium. Targeting was done in the JM8A3.N1 ES cell line, which is derived from a C57BL/6N background and contain an agouti coat color. ES cells were then injected into a C57BL/6 blastocyst by the Rockefeller University Gene Targeting Resource Center. Chimera mice of high chimerism (brown/black in coat color) were selected and bred to C57BL/6 wildtype mice. Offspring were screened for agouti (brown) coat color as an indication of whether the targeted allele was transmitted to the offspring. Several rounds of breeding produced a single brown mouse and southern blotting was used to verify the presence of the RNF126 targeted allele.

Mice containing one copy of the targeted allele (RNF126^ki/+^) are functionally heterozygous because the gene trap construct inserted in intron 1 disrupts translation. The gene trap cassette is flanked by Flipase-recombinase sites (FRT). RNF126^ki/+^ mice were bred to transgenic mice expressing the Flipase recombinase (Jackson Labs # 005703) to reinstate complete translation of the targeted allele. Mice containing a "flipped" version of the targeted allele (RNF126^Fl/+^) were first bred to C57BL/6 mice to cross out the Flipase transgene and then to mb-1 Cre expressing mice ^13^ to generate conditional knockout mice (RNF126^Fl/Fl^mb1^Cre/+^) which were then validated at the DNA, RNA and protein levels as B cell-specific RNF126 conditional knockout mice. RNF126^FL/FL^ mb1^+/+^ were used as wildtype littermate controls in all experiments.

### B Cell Culture Conditions

B Cells (primary and CH12) were cultured in RPMI-1640 + Glutamax (Gibco Life Technologies) supplemented with 10% FBS (BenchMark), 1X Penn Strep (Life Technologies), 55mM b-mercaptoethanol (Life Technologies), 1mM Sodium Pyruvate (Life Technologies), 5mM HEPES (Life Technologies), 1X MEM Non Essential Amino Acids (Life Technologies). HEK 293T cells were cultured in DMEM media (Gibco Life Technologies) supplemented with 10% FBS (BenchMark), 1X Penn Strep (Life Technologies) and 2mM L-glutamine (Life Technologies).

### Viral Transduction of CH12 Cells

HEK 293T cells were transfected with 1 mg of hairpin expressing pLKO.1 lentiviral vectors with 750 ng of psPAX2 and 250ng of pMD2.G packaging vectors with Lipofectamine 2000 reagent (Invitrogen). 24hr post-transfection, media was changed to IMDM (Life Technologies) with 5% FBS (BenchMark). 72hr post-transfection, virus-containing supernatant was collected, filtered through a .45mm filter, and supplemented with polybrene at 8mg/mL concentration. 1X10^6^ CH12 cells were resuspended in viral supernatant and spin-infected at 850g for 2hr at 20C. CH12 cells were removed from the centrifuge and 1mL of culturing medium added.

### shRNA Knockdown Experiments

CH12F3 cells were virally transduced with hairpin expressing vectors against GFP, AID, and RNF126. 48hr post-infection, cells were selected in 1mg/mL of puromycin. Cells were stimulated for CSR with 10 ng/mL IL-4 (Sigma), 1 mg/mL anti-CD40 (eBiosciences) and 1 ng/mL TGFb. CSR to IgA was analyzed by flow cytometry at 48hr post stimulation. Hairpin sequences were as follows:

shGFP: GCAAGCTGACCCTGAAGTTCA,

shAID: CCGGGCGAGATGCATTTCGTATGTTCTCGAGAACATACGAAATGCATCTCGCTTTTTG,

shRNF126-1: CCGGGCTTTGAAATAAATGGACGTTCTCGAGAACGTCCATTTATTTCAAAGCTTTTTG,

shRNF126-2: CCGGGCTCCTCAATCAGTTTGAGAACTCGAGTTCTCAAACTGATTGAGGAGCTTTTTG,

shRNF126-3: CCGGCCCAGTGTGTAAAGAAGACTACTCGAGTAGTCTTCTTTACACACTGGGTTTTTG.

### Splenic B cell Purification and CSR assays

Naïve splenic B cells from 8-12 week old mice were purified by CD43 negative selection on a magnetic column (MACS, Miltenyi Biotec). Purified B cells were plated at a concentration of 0.5 X 10^6^ cells/mL and stimulated with either (1) 10 ng/mL IL-4 (Sigma) and 1 mg/mL anti-CD40 (ebiosciences), (2) 10 ng/mL IL-4 and 25 mg/mL LPS (Sigma) or (3) 25 mg/mL LPS (Sigma). At stated time points post-stimulation (e.g. 72hr, 96hr), cells were collected and FACS analysis was used to determine the percentage of B cells, which have undergone CSR.

### CFSE labeling/proliferation studies

CFSE (Carboxyfluorescein diacetate succinimidyl ester, Invitrogen, CellTrace) was dissolved in DMSO at a final concentration of 5mM. B cells were resuspended at 1X10^6^/mL in PBS/5% FBS. 2mL of CFSE was added for every 1mL of cells to achieve a final CFSE concentration of 10mM. Cells were labeled at room temperature for 10min. The tube was covered with aluminum foil to prevent bleaching. The reaction was quenched by adding an equal volume of FBS and placing the tube on ice for 5min. The tube was filled with PBS/5%FBS and centrifuged at 300xg for 5min at room temperature. Cells were washed two more times in PBS/5%FBS, resuspended in culturing medium supplemented with CSR stimuli and plated. Cells not loaded with CFSE were used as a negative control for FACS analysis.

### Preparation of mammalian cell extracts

Cells were harvested and the pellets washed with 1X PBS. Cells were lysed with RIPA buffer (50mM Tris (pH 8), 150mM NaCl, .5% Na Deoxycholate, .1% SDS, 1% NP-40, .5mM EDTA, 1mM DTT and protease inhibitor cocktail (Roche)). Lysates were normalized with the detergent compatible DC-Assay (Biorad) and probed with anti-RNF126 (Sigma, HPA043050) or anti-AID antibodies (a gift from K. McBride).

### Antibodies Used

Anti-RNF126 (Sigma, HPA043050), CD19-PE (Clone ID3, BD Pharmingen 557399), CD19-APC (Clone ID3, BD Pharmingen 550992), FAS-PE (BD Pharmingen 554258), GL7-FITC (BD Pharmingen 553666), GL7-Alexa Fluor 647 (eBiosciences 51-5902-80), IgG1-PE (Clone A85-1, BD Pharmingen 550083), IgG1-APC (BD Pharmingen 550874), CD43-PE (BD Pharmingen 553271), IgM-PE (BD Pharmingen 553409), IgD FITC (BD Pharmingen 553439), IgA-PE (eBiosciences 12-4204-82).

### Translocation Assays

Live B cells, collected at 72h after CSR stimulation, were the source of genomic DNA that was then analyzed for reciprocal translocations, using previously published assays ^25^.

### Mutation Analysis

Splenic germinal center B cells, or B cells from Peyer’s Patch were FACS-sorted using the surface markers CD19, FAS and GL7 and genomic DNA was prepared. The following regions were amplified with a nested PCR reaction using PFUTurbo (Agilent) and the stated primers:

<u>V186.2 Exon (351 bp amplicon)</u>

5’-TCTTTACAGTTACTGAGCACACAGGAC-3’, 5’GGGTCTAGAGGTGTCCCTAGTCCTTCATGACC-3’

followed by

5’-CAGTAGCAGGCTTGAGGTCTGGAC-3’, 5’GGGTCTAGAGGTGTCCCTAGTCCTTCATGACC-3’),

<u>JH4 Intron (357 bp amplicon)</u>

5’- AGCCTGACATCTGAGGAC-3’, 5’-TAGTGTGGAACATTCCTCAC-3’

followed by

5’-CTGACATCTGAGGACTCTGC-3’, 5’-GCTGTCACAGAGGTGGTCCTG-3’)

<u>V186.2 Upstream Region (295 bp amplicon)</u>

5’-GGCTCTAATGTTACATCCATAGCCTCAAC-3’, 5’GGGTCTAGAGGTGTCCCTAGTCCTTCATGACC-3’

followed by

5’-CAGACAAGATGAGGACTTGGGCTTCAGTATCC-3’, 5’-GTCCAGACCTCAAGCCTGCTACTG-3’).

Amplicons were blunt cloned into the pSC-B vector provided with the StrataClone Blunt PCR Cloning Kit (Agilent) and resulting colonies were sequenced. To align sequences to the consensus and trim vector sequence information external to the PCR product we used BLAT. For each region analyzed, all sequences from one mouse (exlcuding sequences containing indels) were combined and mutation analysis was conducted using SHMTool ^39^. Finally, sequences with indels were analyzed using the MultAlign tool.

